# SARS-CoV-2 spike protein unlikely to bind to integrins via the Arg-Gly-Asp (RGD) motif of the Receptor Binding Domain: evidence from structural analysis and microscale accelerated molecular dynamics

**DOI:** 10.1101/2021.05.24.445335

**Authors:** Houcemeddine Othman, Haifa Ben Messaoud, Oussema Khamessi, Hazem Ben Mabrouk, Kais Ghedira, Avani Bharuthram, Florette Treurnicht, Ikechukwu Achilonu, Yasien Sayed, Najet Srairi-Abid

## Abstract

The Receptor Binding Domain (RBD) of SARS-CoV-2 virus harbors a sequence of Arg-Gly-Asp tripeptide named RGD motif, which has also been identified in extracellular matrix proteins that bind integrins as well as other disintegrins and viruses. Accordingly, integrins have been proposed as host receptors for SARS-CoV-2. The hypothesis was supported by sequence and structural analysis. However, given that the microenvironment of the RGD motif imposes structural hindrance to the protein-protein association, the validity of this hypothesis is still uncertain. Here, we used normal mode analysis, accelerated molecular dynamics microscale simulation, and protein-protein docking to investigate the putative role of RGD motif of SARS-CoV-2 RBD for interacting with integrins. We found, by molecular dynamics, that neither RGD motif nore its microenvironment show any significant conformational shift in the RBD structure. Highly populated clusters were used to run a protein-protein docking against three RGD-binding integrin types, showing no capability of the RBD domain to interact with the RGD binding site. Moreover, the free energy landscape revealed that the RGD conformation within RBD could not acquire an optimal geometry to allow the interaction with integrins. Our results highlighted different structural features of the RGD motif that may prevent its involvement in the interaction with integrins. We, therefore, suggest, in the case where integrins are confirmed to be the direct host receptors for SARS-CoV-2, a possible involvement of other residues to stabilize the interaction.

## 1. Introduction

The molecular mechanism of human infection with SARS-CoV-2 has been studied extensively [47, 35, 28]. Alveolar epithelial cells are thought to be the main target for the virus. Indeed, in pioneering work, Chu *et al.*, [15] studied the tropism of SARS-CoV-2 by inoculating it into 24 cell lines covering seven organs and tracts. They found that the virus most efficiently replicas on lung-type cell lines. Other organs can also be targeted including intestinal tracts, liver, and kidney (*idem*). At the molecular level, the interaction with the host cell involves primarily the homotrimeric spike protein (S protein) expressed on the virus surface. Prior to cell attachment, the spike protein arranges its three Receptor Binding Domains (RBD) in a laying-down configuration, which could help to evade the immune system [10]. Human viruses frequently use mammalian cell surface receptors to attach and to enter host cells [49]. During the interaction process with the host cell, the spike protein switches one of the RBD domains to a standing-up configuration, thus exposing the Receptor Binding Motif (RBM) to the interaction surface of the Angiotensin-Converting Enzyme 2 (ACE2) receptor. ACE2 is widely regarded as the main entry point for the virus to the cellular machinery of the host [52, 43]. However, evidence suggests the possibility of other receptors and co-receptors that might be as relevant as ACE2. The proteomic analysis that helped to establish the interactome map, suggested the putative implication of more than 300 host proteins in the interaction with SARS-CoV-2 [24]. While many of these proteins are expected to be false-positive hits, other studies have pointed out the critical role of specific host proteins and macro-molecules as co-receptors [60], such as neuropilin-1 [13], heparan sulfate [17], sialic acids [45], CD147 [1] and GRP78 [30]. Recently, Sigrist *et al.*[50] have identified an Arg-Gly-Asp (RGD) motif in the sequence of the spike RBD which is found to be exposed at the surface of the interaction domain. This motif was originally identified within the extra-cellular matrix proteins, including fibronectin, fibrinogen, vitronectin, and laminin that mediate cell attachment. Integrins are membrane proteins that act as receptors for these cell adhesion molecules via the RGD motif [27]. Besides, three main integrins expressed on airway epithelial cells were described to play an important role in virus infection [31]. *α*_2_*β*_1_, a collagen and laminin receptor, play a critical role in cell infection by echovirus [21]. Based on these findings, Sigrist *et al.*[50] concluded that integrins can also interact with the spike protein. Several other studies have built on this hypothesis to support the role of integrins as spike protein receptors [36, 18, 7] and to exploit the property for potential therapeutic applications [57]. Moreover, Beddingfield *et al.*[7] showed, by *in vitro* analysis, that the interaction with integrins is a plausible hypothesis. Integrins are heterodimeric receptors that interact favorably with the extracellular molecules by forming a cleft at the protein-protein interface between the beta-propeller and a beta1 domains from the alpha and beta subunits *pmid15378069*. The cleft contains the Metal Ion-Dependent Adhesion Site (MIDAS) harboring an Mg^2+^ ion. Differential expression of *α*_2_*β*_1_, *α*_3_*β*_1_, *α*_4_*β*_1_, *α*_5_*β*_1_, *α*_7_*β*_1_, *α*_6_*β*_4_, *α*_9_*β*_1_, *α_V_ β*_5_, *α_V_ β*_6_, *α_V_ β*_8_ integrins was revealed in human lung cells [6, 12, 55]. Indeed, *α*_2_*β*_1_, *α*_3_*β*_1_, *α*_6_*β*_4_, *α*_9_*β*_1_, *α_V_ β*_5_, *α_V_ β*_6_ and *α_V_ β*_8_ are expressed in airway epithelial cells, which are the main target of coronavirus [46]. Among these, only *α_V_ β*_5_, *α_V_ β*_6_ and *α_V_ β*_8_ can recognize RGD motif while *α*_5_*β*_1_ integrin was not shown to be expressed in healthy epithelial cells [49]. The activity of integrins can be inhibited by disintegrin peptides purified from animals such as snakes, scorpions and insects. The majority of these disintegrins incorporate an RGD motif in their sequences [5, 8, 42, 23, 3]. Most of the arguments about the validity of the RGD motif in SARS-CoV-2 RBD as an interacting segment with integrins are supported by sequence-based and structural-based analysis. However, the microenvironment of RGD imposes a critical steric hindrance that could prevent the RBD from optimally interacting with integrins. To investigate the extent of such effect on the RGD/RBD conformational and binding properties, we conducted a computational study involving microscale accelerated molecular dynamics simulation and protein-protein docking.

## 2. Methods

### 2.1. Structural data

All the structures with complete 3D coordinates of the RBD were explored. They include X-ray crystallography and the cryo-electron microscopy structures. The coordinates of the RBD domain were extracted from the entries of the complete spike protein. In total, we obtained 90 Protein Data Bank (PDB) files (Supplementary data 1).

### 2.2. Normal mode analysis

The normal mode analysis (NMA) approach represents an efficient and powerful tool for predicting and characterizing the large-scale conformational transitions in protein structures around their equilibrium fluctuation. For this study, the Bio3D package in R (version 2.4-1.9000) was utilized to conduct a comparative NMA analysis of a large ensemble of structures [22]. All atoms low-frequency normal modes were calculated under the coarse-grained Elastic Network Model (ENM). Prior to the calculation, structures were aligned to an invariant region of RBD residues. Root Mean Squared Inner Product (RMSIP) was computed from the corresponding eigenvectors of the normal modes to calculate a score quantifying the overlap between modes. The RMSIP was calculated between all the pairs of RBD structures from the collected ensemble of PDB files.

### 2.3. Accelerated molecular dynamics

Accelerated molecular dynamics (aMD) enhances the sampling of a protein conformational space by lowering energy barriers of the energy landscape [26]. A bias term is added to the potential energy *V* (*r*) when the value falls below a certain threshold as follows:

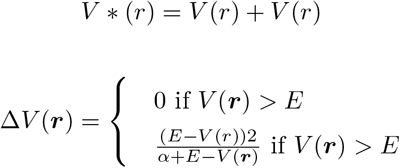

where Δ*V* (****r****) is the bias; *V* (****r****) is the potential energy calculated from the vector of coordinates ****r**** of all the atoms in the system; *E* is the threshold value of the energy, and *α* is the acceleration factor [53]. We used the crystal structure of SARS-CoV2 RBD in complex with H11-D4 antibody (PDB code 6YZ5) at a resolution of 1.8 Å to conduct the simulations. After removing the antibody and the heteroatoms from the structure, we built a Oligomannose-5 glycan (Man5GlcNAc2) type polysaccharide structure and linked it covalently to residue N343 of the RBD (Figure 1)A). The topology of the glycan was identified to be the major form for this amino acid [54]. The system was then neutralized, and TIP3P water molecules were added to a truncated octahedron simulation box where the edges are at a minimum distance of 12 Å for any atom of the solute. Three stages of energy minimization were used to clean the geometry of the atoms and to relax the system. First, we used 5000 steps of steepest descent minimization followed by 15000 steps of conjugate gradient minimization while restraining both water and protein atoms at their initial positions using a force constant of 100 kcal*/*mol*/*A and a non-bonded contact cutoff of 12 Å. We then applied the same minimization series with 400 steps of the steepest descent algorithm and 9600 steps of the conjugate gradient algorithm while applying the constraining force on the protein atoms only. At the final stage, we ran the same cycle and we only lowered the constraining force constant to 0.1 kcal*/*mol*/*Å^2^ applied to the protein atoms. To further relax the system, we applied a heating stage of molecular dynamics by increasing the temperature from 50 K to 300 K while maintaining a force constant of 10 kcal*/*mol*/*Å^2^ on the heavy atoms of the RBD. A Langevin thermostat with a collision frequency of 5 ps^−1^ was applied to control the temperature fluctuation. Following the heating stage, we lifted the constraining forces gradually by an increment of 1 over 11 intervals of 100 ps. The restrained molecular dynamics were run in the NPT ensemble by maintaining the pressure at 1 atm using a relaxation time of 2 ps. The SHAKE method was applied for all the stages of the simulation to constrain the bonds involving hydrogen atoms which allowed an integration time of 100 fs. The Particle Mesh Ewald method was applied to calculate the electrostatic forces. The production phases were run under the NVT conditions. To calculate the different parameters for the aMD simulation, we first run classical molecular dynamics for a total time of 100 ns. From there, we estimated the values of the parameters to calculate the boosting term. The aMD simulation was run in 3 independent replicates for a total time of 1 μs each. An extra boost to the torsional space was added, and the trajectory was constructed by collecting the snapshots at every 10 ps of the running simulation.

**Figure 1:**
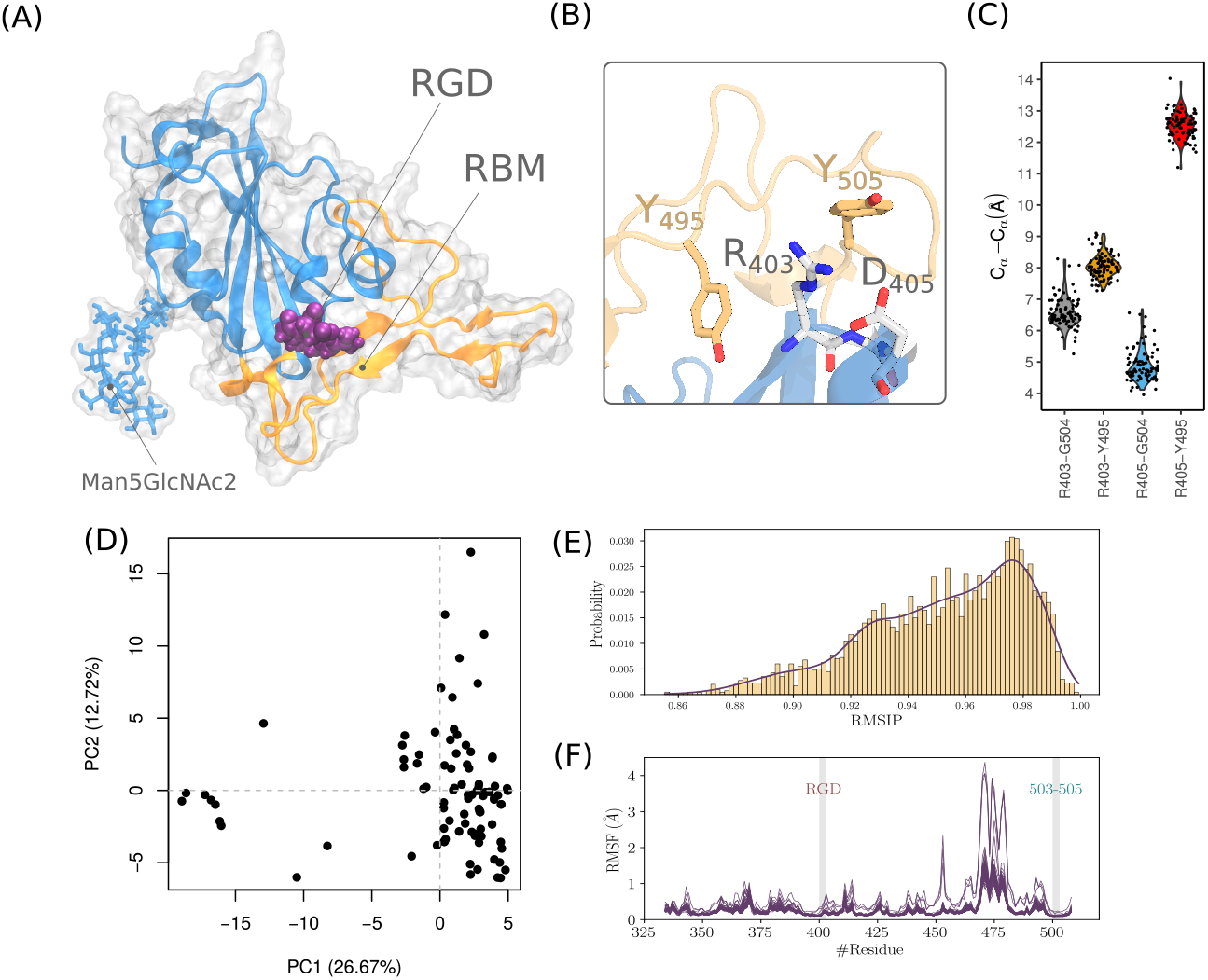
Figure 1. Analysis of the RBD structural ensemble. (A) Structure of RBD showing the RGD motif, the Man5GlcNAc2 polysaccharide and the RBM segment. (B) Arrangement of the RGD motif relative to residue Y505 and Y495. (C) Statistical measurements of distances between RGD residues and D405 and Y495 collected from the ensemble of experimental structures. (D) Projection of RBD structure in the PC1-PC2 subspace of the PCA performed on pre-aligned and superimposed ensemble of structures. (E) RMSIP density plot calculated using the normal modes of each pair of structures of the ensemble. (F) RMSF profile of all the structures in the ensemble computed from all atoms normal mode analysis.

### 2.4. Molecular dynamics data analysis

The crystal structure was set as a reference conformation. Analysis of the molecular dynamics trajectory was made with an in-house python code. Principal Component Analysis (PCA) [19] was calculated for all heavy atoms in the protein, which allowed the detection of dynamical patterns with functional relevance. The translational and rotational related dynamic was first removed by fitting the ensemble of snapshots to the crystal structure of RBD. The low dimension components were calculated to return the corresponding eigenvalues and eigenvectors as well as the projection of the atomic coordinates into the lower-dimensional subspace. Clustering analysis was executed using a hierarchical algorithm embedded in the ‘cpptraj’ analysis tool implemented by AMBER. In this regard an epsilon cutoff of 2 Åwas used. To assess the convergence of the simulation, the cumulative number of clusters (*CNC*) as a function of time and the evolution of informational entropy (*H*) were calculated. The informational entropy is defined by the following formula.

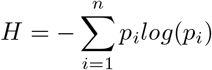

*p_i_* is the probability of the *i^th^* found cluster, as a function of simulation time. To recover the unbiased free energy landscape from the ensemble of conformations sampled by aMD, we reweighted the probability sampling landscape according to the following equation.

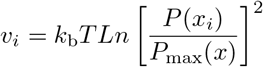

*k_b_* is the Boltzmann constant, T was set to 298 K, *P* (*x_i_*) estimates the probability of a conformational event obtained by binning along the reaction coordinate using the histogram method. The number of bins was set to 50. *P_max_*(*x*) is the maximum probability of the discrete state.

### 2.5. Protein-protein docking

Protein-protein docking was run using the prediction interface of HADDOCK2.2 web server [51]. Integrin structures of *α*_5_*β*_1_, *α*_IIb_*β*_3_, and *α_V_ β*_8_, corresponding to PDB entries 3VI4, 3ZDY and 6UJC respectively, were defined as receptors. The structure of integrins is in a bound state with an RGD binding segment which was removed before running the docking. All residues within a 7 Å distance from the bound RGD in the integrin structure were used to define the active residues of the receptor. Multiple conformations of RBD, compiled from the molecular dynamics simulation, were employed as ligand structures to run the cross-docking. The amino acids of the RGD motif (in position 403-405) were used to define the active residues of the ligand structures. All other parameters of HADDOCK2.2 were kept to their default settings. The structure of the most populated cluster for each docking run was selected for analysis.

## 3. Results

We explored the crystal structure of RBD (PDB code 6YZ5). The RGD motif extends over residues 403-405. R403 is located at the C-terminal end of the fourth *β*-strand of the RBD, while both G404 and D405 are part of its *α*-helix (Figure 1)A). We noticed that only D405 and the guanidinium group of the R403 side chain are solvent-exposed (Figure 1)B). RGD motif shows a considerable kink defined by the main chain atoms and the *C_β_* atoms of R403 and D405. Such configuration leads to the close contact between the RGD motif charged groups with a distance of 4.1 Å. This conformation is different from the optimized configuration of integrin interacting RGDs that adopt an extensive or a slightly kinked configuration [33]. The conformation might be imposed, in part, by the tight interactions with nearby amino acids of the RBD that include Y495 and Y504 (Figure 1)B). Both residues are part of the receptor-binding motif (RBM) with ACE2 [48]. We, therefore, hypothesized that in order to come to an integrin-compatible conformation for RGD, the nearby segments incorporating Y495 and Y504, have to move outwardly relative to the motif. Because the structural properties of the RGD might be imposed by experimental conditions, we tried first to detect major rearrangement within RGD proximity by calculating the distance with Y495 and Y504 from the total set of 90 structures of the RBD. The median distances are 6.4, 8.0, 4.7, and 12.5 Å, corresponding respectively to R403-Y504, R403-Y495, D405-Y504, and D405-Y495 pairs of residues. The distances also show low variability with a maximum difference between the upper and lower values of 2.7 Å noticed for the D405-Y504 pair of residues.

### 3.1. Normal modes analysis

Previous work [4, 9] showed all-atoms elastic network normal mode analysis to be successful in describing the collective dynamics of a wide range of biomolecular systems. We therefore analyzed the ensemble of experimental RBD structures to verify the extent of conformational remodelling that can be adopted and whether it can lead to a better configuration of the RGD atoms in order to be able to interact with integrins. We performed a PCA on the pre-aligned and superimposed ensemble of structures. As shown in the (Figure 1)C), we noticed that variances along PC1 and PC2 can be attributed to few structures of RBD. Most of the projections, however, are located in the lower corner of the PCA plot. We calculated the RMSIP to assess the degree of overlap of the normal modes between the members of the constructed ensemble as proposed in related work [58]. A score of 0.70 is considered a good correspondence, while a score of 0.50 is considered fair [2]. We found that the RMSIP values are ranged from 0.86 to 1 (Figure 1)E) which shows a high level of similarity and agrees with the results from the PCA calculated from the normal modes. We also evaluated the structural deformation adopted by the RBD in terms of Root Mean Square Fluctuation (RMSF) calculated from the projection of the normal modes (Figure 1)F). The structures of the ensemble show an overall similar profile of residue fluctuations in almost all except for some, where increasing flexibility by the amino-acids of the RBM segment is noticed. The profiles revealed limited flexibility for the RGD motif and segment 503-505 with a maximum value of 0.2 Å.

### 3.2. Accelerated molecular dynamics shows local flexibility mainly in the RBM segment but not in RGD microenvironment

Three independent aMD simulations were conducted for a total simulation time of 3 μs. This allows for efficient sampling of the energy landscape for SARS-CoV-2 RBD. The utility of aMD has been previously shown in many macromolecular systems including G-protein coupled receptors, bovine pancreatic trypsin inhibitor, and *α*-1-Antitrypsin [20]. The main goal of this analysis was to identify the most populated conformations that the RBD can take to exert its function of interacting with the host receptors. In the event that the virus binds to integrins via the RGD motif, we would be able to detect a conformational state adapted for such interaction within the set of the sampled aMD snapshots. First, to assess the convergence of the different independent simulations, we calculated the cumulative number of the detected conformational clusters as well as the evolution of Shannon’s entropy (Figure 2)A). We found that, except for one run, all the trajectories show adequate convergence starting from 300 ns in terms of CNCs. The entropy value also converged for all the replicates around 300 ns (Figure 2)B). The coverage of the conformational landscape for RBD was therefore reasonable in the context of our research question. We then verified the conformational drift from the initial structure of RBD for the total *C_α_* atoms, the *C_α_* of the RGD segment, and those of both RGD motif and the 503-505 segment that harbors the Y504 residue (we call this cluster of residues as C1) (Figure 2)C). The latter was included given its proximity to RGD as well as the presumed role that it may play to control the structural properties of the motif. Based on all residue Root Mean Square Deviation (RMSD) values, that can exceed 6 Å, RBD might adopt a significant conformational arrangement. However, the RGD motif does not seem to share this property as the range of RMSDs is less than 0.5 Å. In addition, the C1 residues also did not show a large conformational drift compared to the crystal structure since the corresponding RMSD values are mostly below 2.5 Å. This indeed can also be seen from the RMSF profile of the *C_α_* atoms of RBD (Figure 2)C). The region that shows the largest flexibility corresponds roughly to the RBM residues. The RGD motif shows RMSF values of less than 2 Å while the loop 503-505 has a maximum value of 3.1 Å.

**Figure 2:**
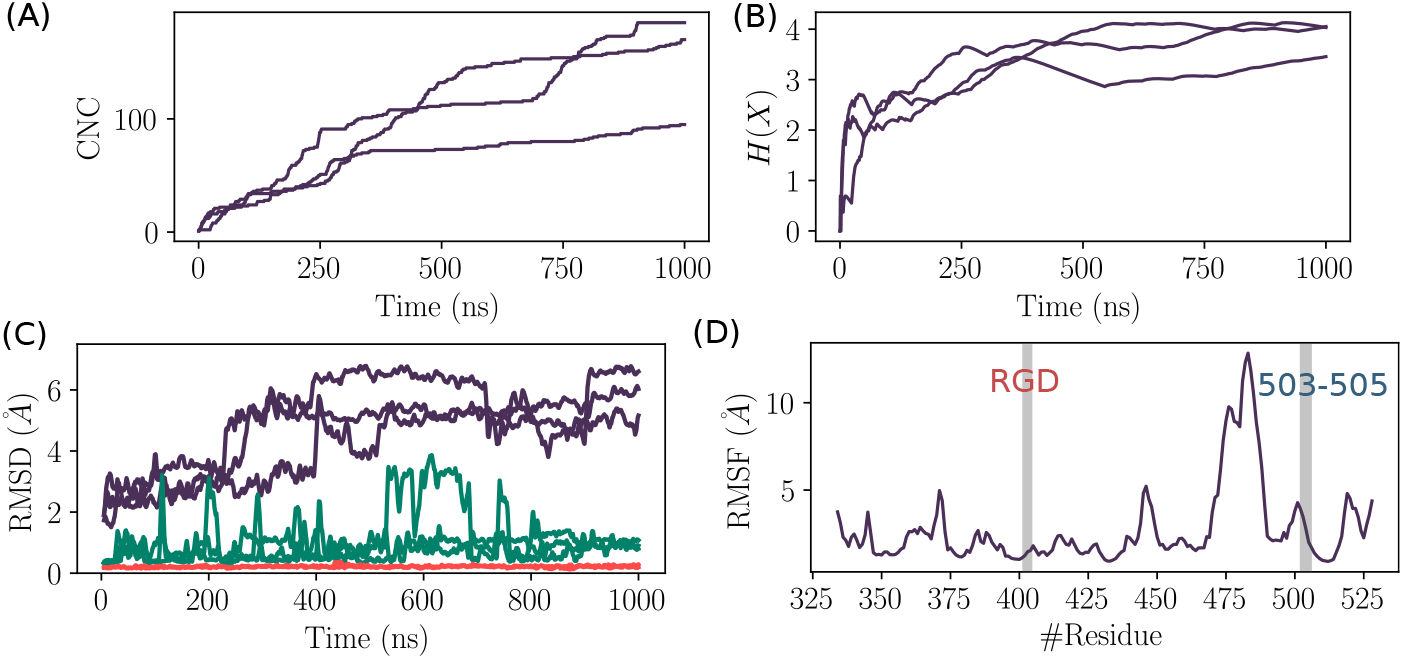
Figure 2. Convergence analysis of aMD and structural deviation of RBD. (A) The cumulative number of clusters as a function of time for the three replicas of aMD trajectories. (B) Evolution of the Shannon’s entropy (*H*(*X*)) for the three replicas of aMD trajectories. (C) Root Mean Square Deviation of RBD structure (Purple), C1 cluster of residues (Green) and the RGD motif (Red). (D) Root Mean Square Fluctuation of RBD residues calculated for the *C*_α_ atoms from the combined aMD trajectories.

### 3.3. PCA and clustering analysis shows no major conformational change in RGD and its microenvironment

We have conducted a principal component analysis using the total set of conformations from the three combined independent trajectories. The protein-heavy atom coordinates were projected onto the subspaces defined by the first and the second components. The aMD simulation was capable of capturing different states of the RBD. We noticed that the structure drifted considerably from the initial crystal structure (red rectangle in Figure 3)A), thus demonstrating the convenient sampling of the RBD phase space that allows ascending the energy barriers. Clustering analysis focused on the clusters showing more than 1% of occupancy. Twenty one major clusters were detected of which the highest-ranked member shows the occupancy of 6.3% (Figure 3)B).

**Figure 3:**
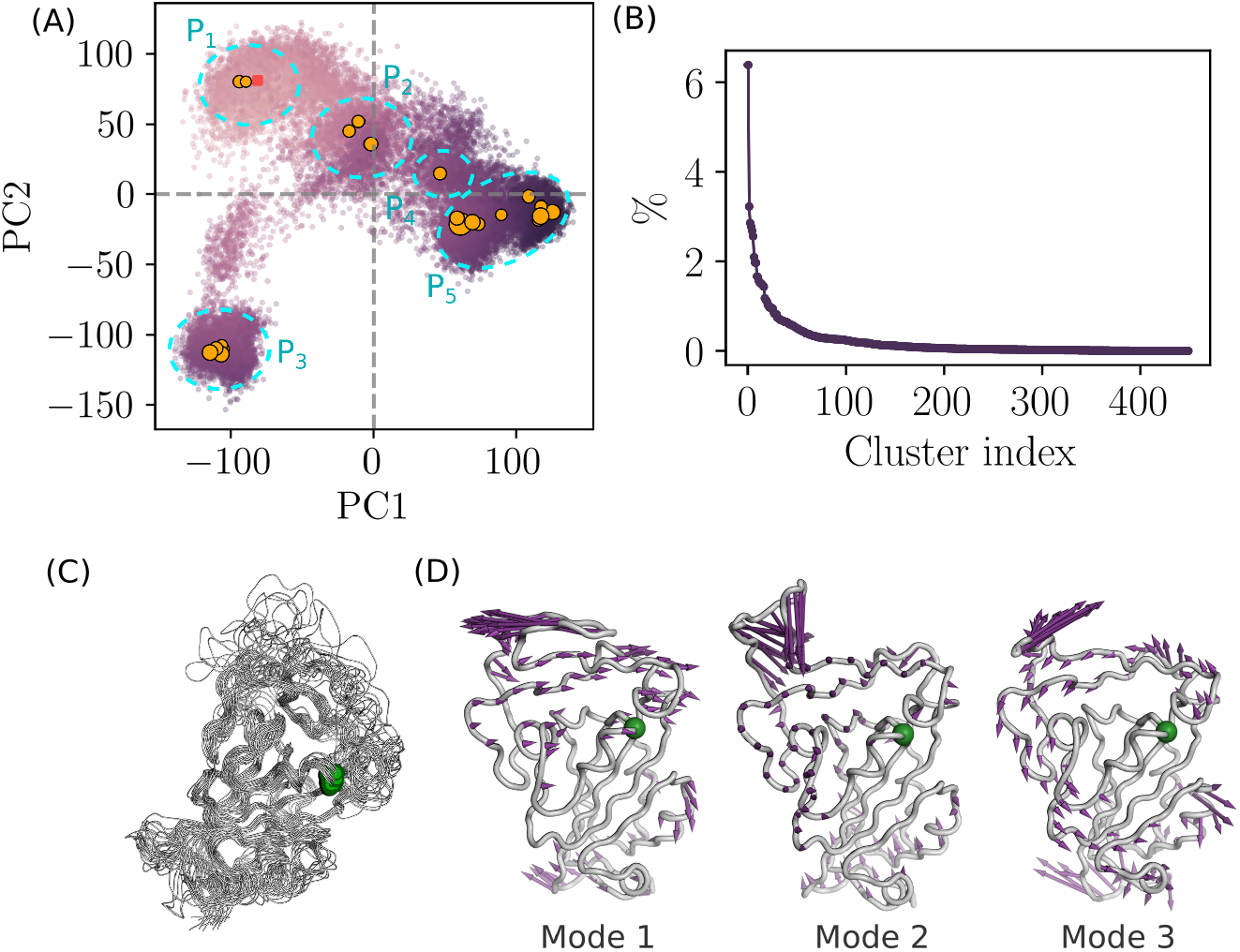
Essential dynamics of RBD from aMD simulation. (A) PCA analysis from the combined replicas. The color of the dots varies as a function of the structural deviation (RMSD) to the crystal structure of RBD. i.e, light purple color indicates lesser deviation and dark purple indicates higher values of RMSD. The square point corresponds to the projection of the crystal structure onto the first and the second subspaces. Orange circles correspond to the centroids of the highly populated clusters and the size of the circles is proportional to the occupancy of the cluster. (B) Occupancy of RBD structural clusters. (C) Structural alignment of the highly populated clusters (occupancy ¿1%). Green spheres indicate the position of the RGD motif. (D) Porcupine plot corresponding to projections of *C*_α_ atoms onto the first three non-rotational and non-translational normal modes.

Essentially, the PCA plot can be subdivided into five different partitions according to the density of the major conformational clusters (Figure 3)A). P1 partition consists of the structures that are close to the bound conformation of RBD. Partitions P2 and P4 correspond to transition states with lower occupancies compared to the other partitions. P3 and P5 correspond to highly populated partitions where the density of the projected atom coordinates is high as shown from the large number of major clusters agglomerated together in the PCA plot. Highly populated partitions, i.e. P1, P3, and P5, may describe the three relevant discrete functional states of RBD corresponding to the bound, up and down states [29]. However, we were unable to verify this, given that the experimental structure of these states lack the atomic details in some RBD segment regions and those at close proximity to the subdomain-1 of the spike protein. Nevertheless, the free energy landscape based on PC1 and PC2, as reaction coordinates established after correcting for the biased sampling of aMD, shows indeed that P2, P3, and P5 correspond to minimum energy wells on the one hand and confirms that P2 and P4 partitions describe transition states on the other hand (Supplementary data 2). Superposition of the representative structures of the highly populated clusters revealed a rigid core of the RBD that harbors the RGD motif of low flexibility (Figure 3)C). Porcupine plots, depicting the direction and the amplitude of motion across the three non-rotational and non-translational normal modes, also highlight the location of the RGD motif within a rigid core of the RBD, characterized by a low amplitude displacement vector (Figure 3)C). Moreover, the RGD motif is rigid in modes 2 and 3, while it moves in the same direction of the segment 503-505 in mode 1.

### 3.4. Favorable geometrical features for the interaction between RGD and integrins are not sampled in the RBD ensemble

Previous research using RGD peptide analogs suggested that extended conformation, spanning the atoms of the aliphatic side chain of Arg and Asp residues as well as the atoms of the main chain of RGD, has to take place to be capable of interacting with integrins [16, 33]. Moreover, the distance between the *C_β_* atoms of Arg and Asp must be within a range of 7 Å to 9 Å. To examine if these properties occurred during aMD simulation, we calculated the angle described by the *C_β_*, *C_α_*, *C_β_* of R403, G404, and D405 residues, respectively, allowing to assess the level of extension (Figure 4)A). We also calculated the distance between the *C_β_* atoms of R403 and D405. 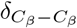 and *θ* describe a wide range of values of 3.6 Å to 9.8 Å and 46° to 172°, respectively (Figure 4)B). However, the data are skewed towards the lower end of the value ranges. Roughly, *θ* has more density in the 46° to 110° range, while the proportion of 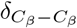 is ranging in higher values of 3.8 Å to 7.7 Å. A strong correlation was also noticed between 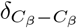 and *θ* with an *R*^2^ value of 0.97 when we fitted the data to a polynomial model. Therefore, we choose the *θ* angle and the RMSD of the C1 cluster of residues as reaction coordinates (Figure 4)C). The FEL has a single highly populated minimum where the values of *θ* roughly span a range of 58° to 83° while the RMSD is low and does not exceed 1.5 Å. Averaging the energy over the binned values of *θ* shows a depth in the energy well of around 3 kcal/mol (Figure 4)D). It also reveals that the more extended *θ* is in the less favorable energy. Indeed the conformation with the lowest energy value shows a significant divergence compared to the states of the RGD motif in its bound form with *α*_5_*β*_1_, *α*_IIb_*β*_3_and *α_V_ β*_8_integrins (Figure 4)E). *θ* and 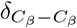 for the lowest energy conformation were measured to 67° and 5.4 Å, respectively. The RGD motif however, clearly adopts an extended conformation in its bound form as revealed by *θ* values of 146°, 173° and 145° and 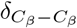 values of 8.9, 9.6 and 8.9 Å for *α*_5_*β*_1_, *α*_IIb_*β*_3_and *α_V_ β*_8_respectively.

**Figure 4:**
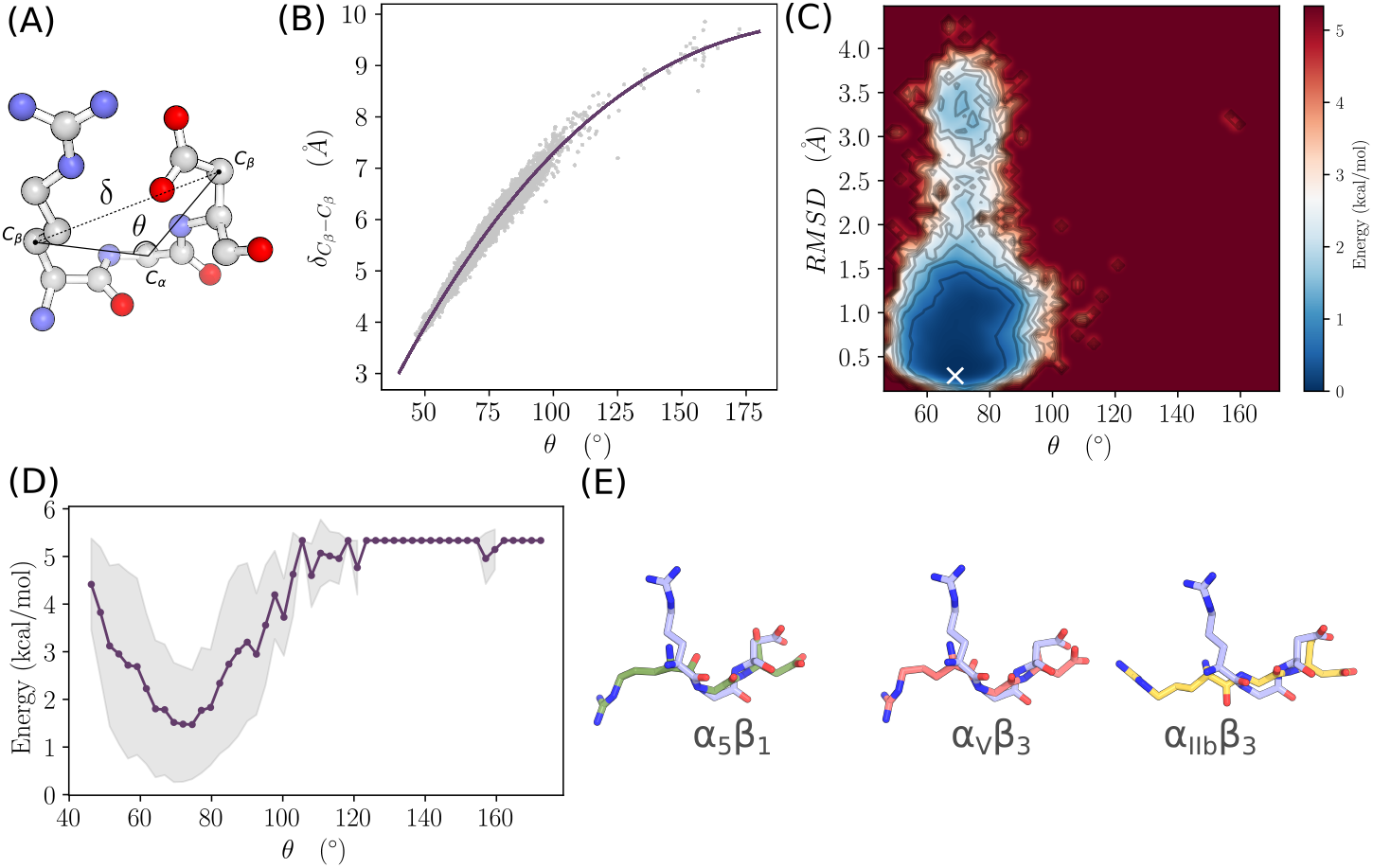
Free energy landscape analysis of the RBD. (A) 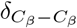 distance and the *θ* angle are indicated on the structure of the RGD segment from RBD. (B) Correlation of 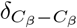 and *θ*. Data were fitted to a polynomial model (*R*^2^ = 0.97). (C) Free energy landscape as a function of *θ* and the RMSD of the C1 residue cluster. The white marker indicates the position of the global minimum. (D) Variation of the energy as a function of *θ*. The gray shading indicates the boundaries defined by the standard deviation of the energy averaged along the reaction coordinate. (E) The RGD structure corresponding to the minimum of energy was fitted and compared to the RGD structure in its bound form with *α*_5_*β*_1_, *α*_IIb_*β*_3_and *α_V_ β*_8_integrins.

### 3.5. Protein-protein docking shows the inability of RGD motif to interact with integrins

We used 22 structures of the highly populated cluster centers obtained from the molecular dynamics simulation to conduct a protein-protein docking. The analysis was conducted by restraining the sampling space to include the RGD motif of RBD and the native binding site on *α*_5_*β*_1_, *α*_IIb_*β*_3_, and *α_V_ β*_8_integrins (Figure 5). These integrins have been chosen mainly for their high-quality crystal structures in a bound state with an RGD motif. Of note, the homology relationship with RGD-binding integrins expressed in airway epithelial cells; namely *α_V_ β*_5_, *α_V_ β*_6_, and *α_V_ β*_8_, is confirmed, implying a conserved 3D fold. Moreover, *α*_IIb_*β*_3_was included to assess the putative binding of SARS-CoV-2 to platelets as suggested by previous studies [61, 34, 59]. Our results show that RBD has not been able to interact favorably with any of the studied integrins. Indeed, RGD motif was not capable of reaching its native binding site in any given structural state.

**Figure 5:**
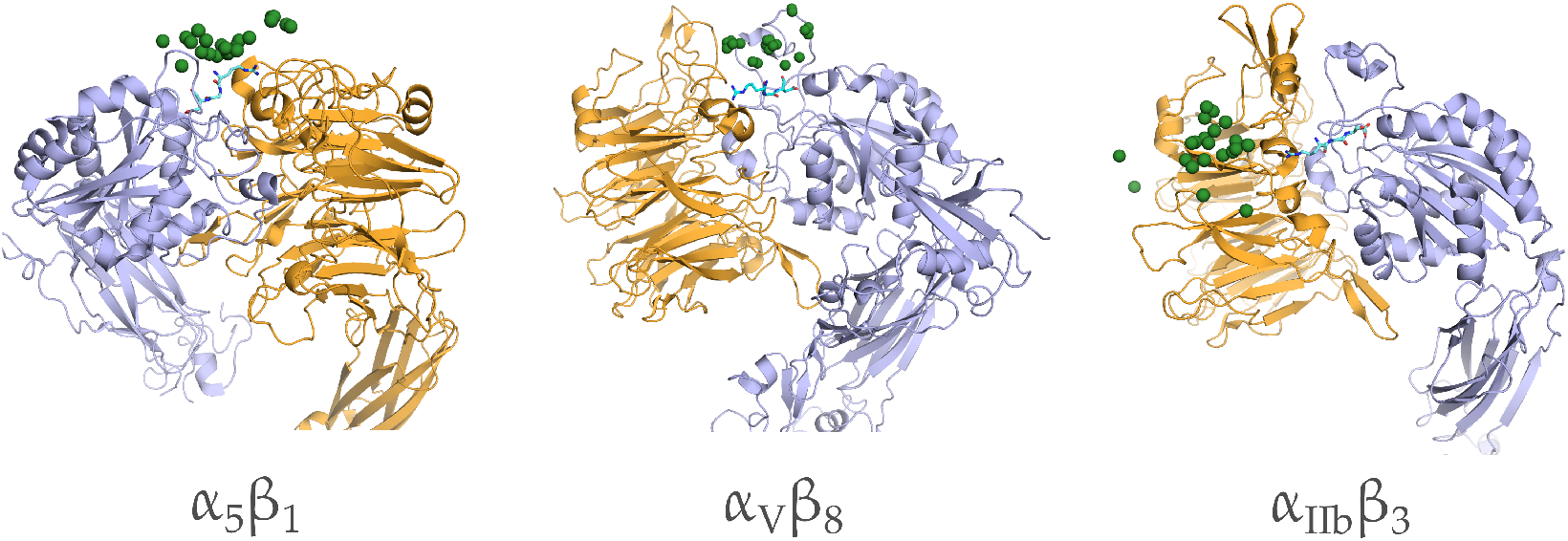
Distribution of the poses of RBD conformations docked to*α*_5_*β*_1_, *α*_IIb_*β*_3_, and *α_V_ β*_8_integrins. The positions of G405 of the RBD motif are shown in green spheres and the native bound configuration of RBD from the crystal structure is shown in cyan sticks.

## 4. Discussion

The optimal interaction of the RGD motif with integrin involves the establishment of a minimal set of contacts with the MIDAS interaction site and the nearby amino acid residues. Experimental structures of RGD in the bound form with integrins show that the motif is laid extensively, crossing the interface cleft between the alpha and beta integrin subunits. The carboxylic and guanidine groups of RGD act as electrostatic clamps with the MIDAS site and the acidic residues of the alpha subunit respectively. However, when we superposed the RGD motif from the RBD domain of SARS-CoV-2 with its corresponding sequence on the cilengitide molecule co-crystallized with the integrin (data not shown), we found that severe clashes persist in this mode of interaction. Following this observation, we hypothesized that the RBD domain must undergo structural adaptation to allow for the favorable interaction with integrins.

The RMSIP distribution demonstrated that the conformational space sampled from the analysis of all the experimental structures are relatively homogeneous, given the observed low variance in the data. Therefore, it is expected that the normal mode properties are linked directly to the conformational behaviour of the RBD. Both normal mode analysis and molecular dynamics simulation are supportive of the relative rigidity of the RGD motif, compared to the RBM amino acids. Therefore, the motif couldn’t undergo a significant structural rearrangement to increase its exposure to the solvent and allow the interaction with integrins. The RGD motif in the structure of different disintegrins, like triflavin, schistatin, echistatin, decorsin and salmosin is located at the tip of a hairpin-like structure that allows an easy fitting with the integration head cleft without steric hindrance [38]. The same type of structure was observed in *α_V_ β*_6_ integrin interacting with the capsid protein VP1 of the foot-and-mouth disease virus (28534487). In the case of SARS-CoV-2 RBD, the RGD motif did not show any structural similarities with disintegrins, and the steric hindrance imposed by the segments close to the motif, seems to be maintained in all the functionally relevant conformational states.

Microscale aMD allowed for an extensive sampling of the conformational phase of RBD where we have detected three highly populated states that could correspond to the bound, up and down configurations of the domain. However, potential integrin-binding conformations were not detected. The free energy landscape also confirmed that the geometrical features of the RGD binding to integrins are unfavorable. Moreover, protein-protein docking showed the inability of all the highly populated conformations to reach the depth of the interaction site of integrins where the electrostatic clamping and the interaction with MIDAS must happen to maintain a stable association.

Most of the former works have relied on sequence conservation and motif detection analysis to conclude on the implication of RGD motif in SARS-CoV-2 RBD in the interaction with integrins [14, 18, 36, 50]. However, few of them have considered the structural features to reinforce or confirm the hypothesis with details, as presented in this study. Indeed, Sigrist *et al.*[50] and Luan *et al.*[36], stated the solvent exposure of RGD as the single argument supporting its involvement in integrin binding, but they did not consider the geometrical features of the motif that must be fulfilled nor the steric hindrance that can be imposed by the surrounding segments. Mèszáros *et al.*[40] and Makowski *et al.*[37] proposed that the surrounding residues of RGD are flexible and, therefore, allow the interaction with integrins. Nevertheless, our results from molecular dynamics simulation and normal mode analysis are congruent in showing that the level of plasticity of these segments is not sufficient to eliminate sterical hindrance that prevents the association with integrins. Moreover, we were not able to detect any hairpin-like structure as observed in disintegrins and VP1 protein of the foot-and-mouth disease virus, despite the extensive sampling of the conformational space. Computational analysis by Dakal [18] concluded that the RGD could bind favorably to *α*_5_*β*_1_ and *α*_5_*β*_6_ integrins. However, in its study the author used only the *β*-propeller head of the alpha subunit for the protein-protein docking, which is not adequate to infer physiological binding properties. On the other hand, Beddingfield *et al.*[7] showed that the protein-protein complex between integrins and S protein, obtained from docking, does not show a favorable fitting in the RGD binding site, which is in agreement with what we have observed from the constrained protein-protein docking analysis.

On the other hand, among integrins identified as expressed in airway epithelial cells, and that could be potential SARS-Cov-2 recepteor, *α*_2_*β*_1_, a collagen and laminin receptor, play a critical role in cell infection by echovirus [21]. This suggests that *α*_2_*β*_1_, which is not an RGD receptor, is unlikely to interact with the motif presented at the surface of the RBD from SARS-CoV-2. The second receptor *α_V_ β*_5_ is well known to be an adenovirus receptor [56] not expressed on the luminal surface [25] which makes it difficult to be involved in the infection by coronavirus. *α_V_ β*_6_, an RGD receptor, was described to be implicated in infection by foot and mouth disease virus [32]. *α_V_ β*_6_ is the only one known to be expressed on the mucosal epithelial cells that are the primary site of infection by respiratory viruses [49]. However, studies using developed antibodies show that *α_V_ β*_6_ is poorly expressed in lung epithelium cells and is constitutively expressed at low levels in uninjured epithelia [11, 55]. Furthermore, the expression pattern of RGD-binding integrins is very differentiated between healthy and unhealthy pulmonary cells. Indeed, many integrins are not seen on healthy adult airway epithelium cells especially *α*_5_*β*_1_ and *α*_9_*β*_1_ [49, 44]. On the other hand, the other expressed RGD dependent integrins have a distinct functional, spatial and chronological expression [44]. *α_V_ β*_5_, *α_V_ β*_6_ and *α_V_ β*_8_ are constitutively expressed at low levels on healthy lung cells [11, 49], recognize many ligands that are not expressed on healthy epithelial basement membranes, and are only involved in cases of lung inflammation and injury [41, 39, 41]. All These informations, consolidated by our above-cited results, emphasize the need for more evidence to confirm the role of integrins in the physiopathology of SARS-CoV-2.

## 5. Conclusion

Based on the evidence provided in this paper, we suggest that the RGD motif from the RBD of SARS-CoV-2 is unlikely to interact with integrins. That, however, does not imply that integrins are not host receptors for the virus. Thus, in light of our results, as well as previous works, the potential interaction of the RGD motif from the RBD of SARS-CoV-2 with integrins should be revised extensively. Consequently, potential involvement of other segments belonging to the spike protein, is more likely to take place if integrins are confirmed to be host receptors for SARS-CoV-2.

## Acknowledgments

The authors acknowledge the Centre for High Performance Computing (CHPC), South Africa, for providing computational resources to this research project. We thank Dr. Fatma Z. Guerfali from Institut Pasteur de Tunis for providing critical reviews of the paper. The present work was partially supported by the European project PHINDaccess: Strengthening Omics data analysis capacities in pathogen-host interaction (Grant Agreement ID: 811034).

## References

[1] J. A. Aguiar, B. J. Tremblay, M. J. Mansfield, O. Woody, B. Lobb, A. Banerjee, A. Chandiramohan, N. Tiessen, Q. Cao, A. Dvorkin-Gheva, S. Revill, M. S. Miller, C. Carlsten, L. Organ, C. Joseph, A. John, P. Hanson, R. C. Austin, B. M. McManus, G. Jenkins, K. Mossman, K. Ask, A. C. Doxey, and J. A. Hirota. Gene expression and in situ protein profiling of candidate SARS-CoV-2 receptors in human airway epithelial cells and lung tissue. Eur Respir J, 56(3), 09 2020.

[2] A. Amadei, M. A. Ceruso, and A. Di Nola. On the convergence of the conformational coordinates basis set obtained by the essential dynamics analysis of proteins’ molecular dynamics simulations. Proteins, 36(4):419–424, Sep 1999.

[3] T. C. Assumpcao, J. M. Ribeiro, and I. M. Francischetti. Disintegrins from hematophagous sources. Toxins (Basel), 4(5):296–322, 05 2012.

[4] I. Bahar, T. R. Lezon, A. Bakan, and I. H. Shrivastava. Normal mode analysis of biomolecular structures: functional mechanisms of membrane proteins. Chem Rev, 110(3):1463–1497, Mar 2010.

[5] A. Bazaa, P. Juárez, N. Marrakchi, Z. Bel Lasfer, M. El Ayeb, R. A. Harrison, J. J. Calvete, and L. Sanz. Loss of introns along the evolutionary diversification pathway of snake venom disintegrins evidenced by sequence analysis of genomic DNA from Macrovipera lebetina transmediterranea and Echis ocellatus. J Mol Evol, 64(2):261–271, Feb 2007.

[6] S. Bazan-Socha, A. Bukiej, C. Marcinkiewicz, and J. Musial. Integrins in pulmonary inflammatory diseases. Curr Pharm Des, 11(7):893–901, 2005.

[7] B. J. Beddingfield, N. Iwanaga, P. P. Chapagain, W. Zheng, C. J. Roy, T. Y. Hu, J. K. Kolls, and G. J. Bix. The Integrin Binding Peptide, ATN-161, as a Novel Therapy for SARS-CoV-2 Infection. JACC Basic Transl Sci, 6(1):1–8, Jan 2021.

[8] H. Ben-Mabrouk, R. Zouari-Kessentini, F. Montassar, Z. A. Koubaa, E. Messaadi, X. Guillonneau, M. ElAyeb, N. Srairi-Abid, J. Luis, O. Micheau, and N. Marrakchi. CC5 and CC8, two homologous disintegrins from Cerastes cerastes venom, inhibit in vitro and ex vivo angiogenesis. Int J Biol Macromol, 86:670–680, May 2016.

[9] A. Bende. Hydrogen bonding in the urea dimers and adenine–thymine dna base pair: anharmonic effects in the intermolecular h-bond and intramolecular h-stretching vibrations. Theoretical Chemistry Accounts, 125(3):253–268, 2010.

[10] J. D. Berry, S. Jones, M. A. Drebot, A. Andonov, M. Sabara, X. Y. Yuan, H. Weingartl, L. Fernando, P. Marszal, J. Gren, et al. Development and characterisation of neutralising monoclonal antibody to the sars-coronavirus. Journal of virological methods, 120(1):87–96, 2004.

[11] J. M. Breuss, J. Gallo, H. M. DeLisser, I. V. Klimanskaya, H. G. Folkesson, J. F. Pittet, S. L. Nishimura, K. Aldape, D. V. Landers, and W. Carpenter. Expression of the beta 6 integrin subunit in development, neoplasia and tissue repair suggests a role in epithelial remodeling. J Cell Sci, 108 (Pt 6):2241–2251, Jun 1995.

[12] S. Cambier, D. Z. Mu, D. O’Connell, K. Boylen, W. Travis, W. H. Liu, V. C. Broaddus, and S. L. Nishimura. A role for the integrin alphavbeta8 in the negative regulation of epithelial cell growth. Cancer Res, 60(24):7084–7093, Dec 2000.

[13] L. Cantuti-Castelvetri, R. Ojha, L. D. Pedro, M. Djannatian, J. Franz, S. Kuivanen, K. Kallio, T. Kaya, M. Anastasina, T. Smura, L. Levanov, L. Szirovicza, A. Tobi, H. Kallio-Kokko, P. Österlund, M. Joensuu, F. A. Meunier, S. Butcher, M. S. Winkler, B. Mollenhauer, A. Helenius, O. Gokce, T. Teesalu, J. Hepojoki, O. Vapalahti, C. Stadelmann, G. Balistreri, and M. Simons. Neuropilin-1 facilitates sars-cov-2 cell entry and provides a possible pathway into the central nervous system. bioRxiv, 2020.

[14] I. Carvacho and M. Piesche. RGD-binding integrins and TGF-*β* in SARS-CoV-2 infections - novel targets to treat COVID-19 patients? Clin Transl Immunology, 10(3):e1240, 2021.

[15] H. Chu, J. F. Chan, T. T. Yuen, H. Shuai, S. Yuan, Y. Wang, B. Hu, C. C. Yip, J. O. Tsang, X. Huang, Y. Chai, D. Yang, Y. Hou, K. K. Chik, X. Zhang, A. Y. Fung, H. W. Tsoi, J. P. Cai, W. M. Chan, J. D. Ip, A. W. Chu, J. Zhou, D. C. Lung, K. H. Kok, K. K. To, O. T. Tsang, K. H. Chan, and K. Y. Yuen. Comparative tropism, replication kinetics, and cell damage profiling of SARS-CoV-2 and SARS-CoV with implications for clinical manifestations, transmissibility, and laboratory studies of COVID-19: an observational study. Lancet Microbe, 1(1):e14–e23, May 2020.

[16] M. Civera, D. Arosio, F. Bonato, L. Manzoni, L. Pignataro, S. Zanella, C. Gennari, U. Piarulli, and L. Belvisi. Investigating the Interaction of Cyclic RGD Peptidomimetics with AlphaVBeta6 Integrin by Biochemical and Molecular Docking Studies. Cancers (Basel), 9(10), Sep 2017.

[17] T. M. Clausen, D. R. Sandoval, C. B. Spliid, J. Pihl, C. D. Painter, B. E. Thacker, C. A. Glass, A. Narayanan, S. A. Majowicz, Y. Zhang, J. L. Torres, G. J. Golden, R. Porell, A. F. Garretson, L. Laubach, J. Feldman, X. Yin, Y. Pu, B. Hauser, T. M. Caradonna, B. P. Kellman, C. Martino, P. L. S. M. Gordts, S. L. Leibel, S. K. Chanda, A. G. Schmidt, K. Godula, J. Jose, K. D. Corbett, B. Ward, A. F. Carlin, and J. D. Esko. SARS-CoV-2 Infection Depends on Cellular Heparan Sulfate and ACE2. bioRxiv, Jul 2020.

[18] T. C. Dakal. SARS-CoV-2 attachment to host cells is possibly mediated via RGD-integrin interaction in a calcium-dependent manner and suggests pulmonary EDTA chelation therapy as a novel treatment for COVID 19. Immunobiology, 226(1):152021, 01 2021.

[19] C. C. David and D. J. Jacobs. Principal component analysis: a method for determining the essential dynamics of proteins. Methods Mol Biol, 1084:193–226, 2014.

[20] L. Duan, X. Guo, Y. Cong, G. Feng, Y. Li, and J. Z. Zhang. Accelerated molecular dynamics simulation for helical proteins folding in explicit water. Frontiers in chemistry, 7:540, 2019.

[21] R. Eisner. Finding out how a viral hitchhiker snags a ride. Science, 255(5052):1647, Mar 1992.

[22] E. Fuglebakk, J. Echave, and N. Reuter. Measuring and comparing structural fluctuation patterns in large protein datasets. Bioinformatics, 28(19):2431–2440, Oct 2012.

[23] A. Gasmi, N. Srairi, S. Guermazi, H. Dekhil, H. Dkhil, H. Karoui, and M. El Ayeb. Amino acid structure and characterization of a heterodimeric disintegrin from Vipera lebetina venom. Biochim Biophys Acta, 1547(1):51–56, May 2001.

[24] D. E. Gordon, G. M. Jang, M. Bouhaddou, J. Xu, K. Obernier, K. M. White, M. J. O’Meara, V. V. Rezelj, J. Z. Guo, D. L. Swaney, T. A. Tummino, R. Hüttenhain, R. M. Kaake, A. L. Richards, A. Tutuncuoglu, H. Foussard, J. Batra, K. Haas, M. Modak, M. Kim, P. Haas, B. J. Polacco, H. Braberg, J. M. Fabius, M. Eckhardt, M. Soucheray, M. J. Bennett, M. Cakir, M. J. McGregor, Q. Li, B. Meyer, F. Roesch, T. Vallet, A. Mac Kain, L. Miorin, E. Moreno, Z. Z. C. Naing, Y. Zhou, S. Peng, Y. Shi, Z. Zhang, W. Shen, I. T. Kirby, J. E. Melnyk, J. S. Chorba, K. Lou, S. A. Dai, I. Barrio-Hernandez, D. Memon, C. Hernandez-Armenta, J. Lyu, C. J. P. Mathy, T. Perica, K. B. Pilla, S. J. Ganesan, D. J. Saltzberg, R. Rakesh, X. Liu, S. B. Rosenthal, L. Calviello, S. Venkataramanan, J. Liboy-Lugo, Y. Lin, X. P. Huang, Y. Liu, S. A. Wankowicz, M. Bohn, M. Safari, F. S. Ugur, C. Koh, N. S. Savar, Q. D. Tran, D. Shengjuler, S. J. Fletcher, M. C. O’Neal, Y. Cai, J. C. J. Chang, D. J. Broadhurst, S. Klippsten, P. P. Sharp, N. A. Wenzell, D. Kuzuoglu-Ozturk, H. Y. Wang, R. Trenker, J. M. Young, D. A. Cavero, J. Hiatt, T. L. Roth, U. Rathore, A. Subramanian, J. Noack, M. Hubert, R. M. Stroud, A. D. Frankel, O. S. Rosenberg, K. A. Verba, D. A. Agard, M. Ott, M. Emerman, N. Jura, M. von Zastrow, E. Verdin, A. Ashworth, O. Schwartz, C. d’Enfert, S. Mukherjee, M. Jacobson, H. S. Malik, D. G. Fujimori, T. Ideker, C. S. Craik, S. N. Floor, J. S. Fraser, J. D. Gross, A. Sali, B. L. Roth, D. Ruggero, J. Taunton, T. Kortemme, P. Beltrao, M. Vignuzzi, A. García-Sastre, K. M. Shokat, B. K. Shoichet, and N. J. Krogan. A SARS-CoV-2 protein interaction map reveals targets for drug repurposing. Nature, 583(7816):459–468, 07 2020.

[25] B. R. Grubb, R. J. Pickles, H. Ye, J. R. Yankaskas, R. N. Vick, J. F. Engelhardt, J. M. Wilson, L. G. Johnson, and R. C. Boucher. Inefficient gene transfer by adenovirus vector to cystic fibrosis airway epithelia of mice and humans. Nature, 371(6500):802–806, Oct 1994.

[26] D. Hamelberg, J. Mongan, and J. A. McCammon. Accelerated molecular dynamics: a promising and efficient simulation method for biomolecules. J Chem Phys, 120(24):11919–11929, Jun 2004.

[27] H. Hamidi, M. Pietilä, and J. Ivaska. The complexity of integrins in cancer and new scopes for therapeutic targeting. Br J Cancer, 115(9):1017–1023, Oct 2016.

[28] A. G. Harrison, T. Lin, and P. Wang. Mechanisms of SARS-CoV-2 Transmission and Pathogenesis. Trends Immunol, 41(12):1100–1115, 12 2020.

[29] R. Henderson, R. J. Edwards, K. Mansouri, K. Janowska, V. Stalls, S. M. C. Gobeil, M. Kopp, D. Li, R. Parks, A. L. Hsu, M. J. Borgnia, B. F. Haynes, and P. Acharya. Controlling the SARS-CoV-2 spike glycoprotein conformation. Nat Struct Mol Biol, 27(10):925–933, 10 2020.

[30] I. M. Ibrahim, D. H. Abdelmalek, M. E. Elshahat, and A. A. Elfiky. COVID-19 spike-host cell receptor GRP78 binding site prediction. J Infect, 80(5):554–562, 05 2020.

[31] R. R. Isberg and G. Tran Van Nhieu. Binding and internalization of microorganisms by integrin receptors. Trends Microbiol, 2(1):10–14, Jan 1994.

[32] T. Jackson, D. Sheppard, M. Denyer, W. Blakemore, and A. M. King. The epithelial integrin alphavbeta6 is a receptor for foot-and-mouth disease virus. J Virol, 74(11):4949–4956, Jun 2000.

[33] T. G. Kapp, F. Rechenmacher, S. Neubauer, O. V. Maltsev, E. A. Cavalcanti-Adam, R. Zarka, U. Reuning, J. Notni, H. J. Wester, C. Mas-Moruno, J. Spatz, B. Geiger, and H. Kessler. A Comprehensive Evaluation of the Activity and Selectivity Profile of Ligands for RGD-binding Integrins. Sci Rep, 7:39805, 01 2017.

[34] M. Koupenova and J. E. Freedman. Platelets and COVID-19: Inflammation, Hyperactivation and Additional Questions. Circ Res, 127(11):1419–1421, 11 2020.

[35] S. Li, S. Li, C. Disoma, R. Zheng, M. Zhou, A. Razzaq, P. Liu, Y. Zhou, Z. Dong, A. Du, et al. Sars-cov-2: Mechanism of infection and emerging technologies for future prospects. Reviews in Medical Virology, 31(2):e2168, 2021.

[36] J. Luan, Y. Lu, S. Gao, and L. Zhang. A potential inhibitory role for integrin in the receptor targeting of SARS-CoV-2. J Infect, 81(2):318–356, 08 2020.

[37] L. Makowski, W. Olson-Sidford, and J. W-Weisel. Biological and Clinical Consequences of Integrin Binding via a Rogue RGD Motif in the SARS CoV-2 Spike Protein. Viruses, 13(2), Jan 2021.

[38] T. Matsui, J. Hamako, and K. Titani. Structure and function of snake venom proteins affecting platelet plug formation. Toxins (Basel), 2(1):10–23, 01 2010.

[39] D. Mu, S. Cambier, L. Fjellbirkeland, J. L. Baron, J. S. Munger, H. Kawakatsu, D. Sheppard, C. Broaddus, and S. L. Nishimura. The integrin alpha(v)beta8 mediates epithelial homeostasis through MT1-MMP-dependent activation of TGF-beta1. J Cell Biol, 157(3):493–507, Apr 2002.

[40] B. Mészáros, H. Sámano-Sánchez, J. Alvarado-Valverde, J. Čalyševa, E. Martínez-Pérez, R. Alves, D. C. Shields, M. Kumar, F. Rippmann, L. B. Chemes, and T. J. Gibson. Short linear motif candidates in the cell entry system used by SARS-CoV-2 and their potential therapeutic implications. Sci Signal, 14(665), 01 2021.

[41] S. L. Nishimura, D. Sheppard, and R. Pytela. Integrin alpha v beta 8. Interaction with vitronectin and functional divergence of the beta 8 cytoplasmic domain. J Biol Chem, 269(46):28708–28715, Nov 1994.

[42] K. Z. Olfa, L. José, D. Salma, B. Amine, S. A. Najet, A. Nicolas, L. Maxime, Z. Raoudha, M. Kamel, M. Jacques, S. Jean-Marc, e. l. A. Mohamed, and M. Naziha. Lebestatin, a disintegrin from Macrovipera venom, inhibits integrin-mediated cell adhesion, migration and angiogenesis. Lab Invest, 85(12):1507–1516, Dec 2005.

[43] H. Othman, Z. Bouslama, J.-T. Brandenburg, J. Da Rocha, Y. Hamdi, K. Ghedira, N. Srairi-Abid, and S. Hazelhurst. Interaction of the spike protein rbd from sars-cov-2 with ace2: Similarity with sars-cov, hot-spot analysis and effect of the receptor polymorphism. Biochemical and biophysical research communications, 527(3):702–708, 2020.

[44] J. M. Pilewski, J. D. Latoche, S. M. Arcasoy, and S. M. Albelda. Expression of integrin cell adhesion receptors during human airway epithelial repair in vivo. Am J Physiol, 273(1 Pt 1):L256–263, Jul 1997.

[45] E. Qing, M. Hantak, S. Perlman, and T. Gallagher. Distinct Roles for Sialoside and Protein Receptors in Coronavirus Infection. mBio, 11(1), 02 2020.

[46] N. G. Ravindra, M. M. Alfajaro, V. Gasque, V. Habet, J. Wei, R. B. Filler, N. C. Huston, H. Wan, K. Szigeti-Buck, B. Wang, G. Wang, R. R. Montgomery, S. C. Eisenbarth, A. Williams, A. M. Pyle, A. Iwasaki, T. L. Horvath, E. F. Foxman, R. W. Pierce, D. van Dijk, and C. B. Wilen. Single-cell longitudinal analysis of SARS-CoV-2 infection in human bronchial epithelial cells. bioRxiv, May 2020.

[47] J. Shang, Y. Wan, C. Luo, G. Ye, Q. Geng, A. Auerbach, and F. Li. Cell entry mechanisms of SARS-CoV-2. Proc Natl Acad Sci U S A, 117(21):11727–11734, 05 2020.

[48] J. Shang, G. Ye, K. Shi, Y. Wan, C. Luo, H. Aihara, Q. Geng, A. Auerbach, and F. Li. Structural basis of receptor recognition by SARS-CoV-2. Nature, 581(7807):221–224, 05 2020.

[49] D. Sheppard. Functions of pulmonary epithelial integrins: from development to disease. Physiol Rev, 83(3):673–686, Jul 2003.

[50] C. J. Sigrist, A. Bridge, and P. Le Mercier. A potential role for integrins in host cell entry by SARS-CoV-2. Antiviral Res, 177:104759, 05 2020.

[51] G. C. P. van Zundert, J. P. G. L. M. Rodrigues, M. Trellet, C. Schmitz, P. L. Kastritis, E. Karaca, A. S. J. Melquiond, M. van Dijk, S. J. de Vries, and A. M. J. J. Bonvin. The HADDOCK2.2 Web Server: User-Friendly Integrative Modeling of Biomolecular Complexes. J Mol Biol, 428(4):720–725, Feb 2016.

[52] Y. Wan, J. Shang, R. Graham, R. S. Baric, and F. Li. Receptor recognition by the novel coronavirus from wuhan: an analysis based on decade-long structural studies of sars coronavirus. Journal of Virology, 94(7), 2020.

[53] Y. Wang, C. B. Harrison, K. Schulten, and J. A. McCammon. Implementation of Accelerated Molecular Dynamics in NAMD. Comput Sci Discov, 4(1), 2011.

[54] Y. Watanabe, J. D. Allen, D. Wrapp, J. S. McLellan, and M. Crispin. Site-specific analysis of the sars-cov-2 glycan shield. BioRxiv, 2020.

[55] A. Weinacker, R. Ferrando, M. Elliott, J. Hogg, J. Balmes, and D. Sheppard. Distribution of integrins alpha v beta 6 and alpha 9 beta 1 and their known ligands, fibronectin and tenascin, in human airways. Am J Respir Cell Mol Biol, 12(5):547–556, May 1995.

[56] T. J. Wickham, E. J. Filardo, D. A. Cheresh, and G. R. Nemerow. Integrin alpha v beta 5 selectively promotes adenovirus mediated cell membrane permeabilization. J Cell Biol, 127(1):257–264, Oct 1994.

[57] S. Yan, H. Sun, X. Bu, and G. Wan. New Strategy for COVID-19: An Evolutionary Role for RGD Motif in SARS-CoV-2 and Potential Inhibitors for Virus Infection. Front Pharmacol, 11:912, 2020.

[58] X.-Q. Yao, L. Skjærven, and B. J. Grant. Rapid characterization of allosteric networks with ensemble normal mode analysis. The Journal of Physical Chemistry B, 120(33):8276–8288, 2016.

[59] Y. Zaid, F. Puhm, I. Allaeys, A. Naya, M. Oudghiri, L. Khalki, Y. Limami, N. Zaid, K. Sadki, R. Ben El Haj, et al. Platelets can associate with sars-cov-2 rna and are hyperactivated in covid-19. Circulation research, 127(11):1404–1418, 2020.

[60] N. Zamorano Cuervo and N. Grandvaux. ACE2: Evidence of role as entry receptor for SARS-CoV-2 and implications in comorbidities. Elife, 9, 11 2020.

[61] S. Zhang, Y. Liu, X. Wang, L. Yang, H. Li, Y. Wang, M. Liu, X. Zhao, Y. Xie, Y. Yang, S. Zhang, Z. Fan, J. Dong, Z. Yuan, Z. Ding, Y. Zhang, and L. Hu. SARS-CoV-2 binds platelet ACE2 to enhance thrombosis in COVID-19. J Hematol Oncol, 13(1):120, 09 2020.

