## Supplemental data for "SARS-CoV-2 spike protein unlikely to bind to integrins via the Arg-Gly-Asp (RGD) motif of the Receptor Binding Domain: evidence from structural analysis and microscale accelerated molecular dynamics"

[a]*Sydney Brenner Institute for Molecular Bioscience, Faculty of Health Sciences, University of the Witwatersrand, Johannesburg, South Africa.*
[b]*National Gene Bank of Tunisia, Boulevard du Leader Yesser Arafet, Tunis, Tunisia.*
[c]*Université de Tunis El Manar, Institut Pasteur de Tunis, LR11IPT08 Venins et Biomolecules Therapeutiques, 1002, Tunis, Tunisie.*
[d]*Loboratory of Biomolecules, Venoms and Theranostic Applications, LR20IPT01, Institut Pasteur de Tunis, 13, Place Pasteur. University of Tunis El Manar, Tunis,Tunisia.*
[e]*Laboratory of Bioinformatics, Biomathematics and Biostatistics (BIMS), Institut Pasteur de Tunis (IPT), 13, Place Pasteur BP 74. University of Tunis El Manar, Tunis,Tunisia.*
[f]*Department of Virology, National Health Laboratory Services and the School of Pathology, University of the Witwatersrand, Johannesburg, South Africa.*
[g]*Protein Structure-Function Research Unit, School of Molecular and Cell Biology, University of Witwatersrand, Johannesburg, South Africa.*

### 1. Supplementary material 1

List of structures included in the normal mode analysis:
6LZG, 6M0J, 6M17, 6W41, 6XC2, 6XC3, 6XC4, 6XC7, 6XCN, 6XDG, 6XE1, 6XEY, 6XF5, 6XF6, 6XKP, 6XLU, 6XM0, 6XM3, 6XM4, 6XM5, 6YLA, 6YM0, 6YOR, 6YZ5, 6Z2M, 6Z43, 6Z97, 6ZB4, 6ZB5, 6ZCZ, 6ZDG, 6ZDH, 6ZER, 6ZFO, 6ZGE, 6ZGG, 6ZGI, 6ZHD, 6ZOX, 6ZOY, 6ZOZ, 6ZP0, 6ZP1, 6ZP2, 6ZXN, 7A29, 7A4N, 7A93, 7A94, 7A95, 7A96, 7A97, 7A98, 7BWJ, 7BZ5, 7C01, 7C8D, 7C8V, 7C8W, 7CAI, 7CAK, 7CAN, 7CH4, 7CH5, 7CHB, 7CHC, 7CHE, 7CHF, 7CHH, 7JJI, 7JMO, 7JMW, 7JVC, 7JW0, 7JX3, 7JZL, 7JZM, 7JZN, 7JZU, 7K43, 7K45, 7K4N, 7K8S, 7K8T, 7K8U, 7K8W, 7K8X, 7K8Y, 7K90, 7K9Z.

[1]These authors contributed equally to the work.

**2. Supplementary material 2**

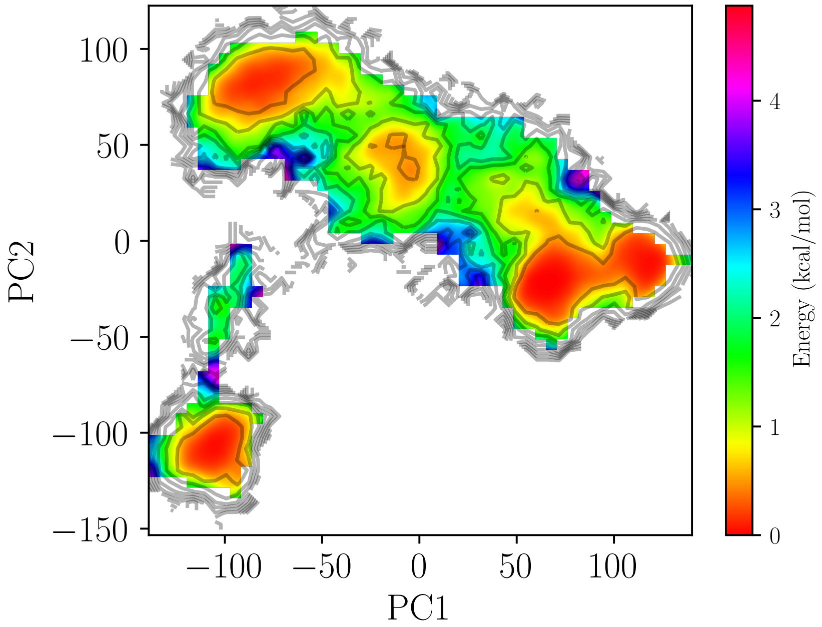


Figure 1: Free energy landscape of RBD from SARS-CoV-2 constructed from PC1 and PC2 as reaction coordinates.
